# Quantifying Forest Biomass and Genetic Contribution using Light Detection and Ranging

**DOI:** 10.1101/2024.07.03.601985

**Authors:** Haruka Sano, Naoko Miura, Minoru Inamori, Yamato Unno, Wei Guo, Sachiko Isobe, Kazutaka Kusunoki, Hiroyoshi Iwata

**Affiliations:** Graduate School of Agricultural and Life Sciences, The University of Tokyo; SUMITOMO FORESTRY CO., LTD; Kazusa DNA Research Institute

## Abstract

The growing focus on the role of forests in carbon sequestration highlights the importance of accurately and efficiently measuring biophysical traits, such as diameter at breast height (DBH) and tree height. Understanding genetic contributions to trait variation is crucial for enhancing carbon storage through genetic improvement of forest trees. Light detection and ranging (LiDAR) has been used to estimate DBH and tree height; however, few studies have explored the heritability of these traits or assessed the accuracy of biomass increment selections based on these traits. Therefore, this study aimed to leverage LiDAR to measure DBH and tree height, estimate tree heritability, and evaluate the accuracy of timber volume selections based on these traits using 60-year-old larch as the study material. Unmanned aerial vehicle (UAV) and backpack LiDAR were compared against hand-measured values. The accuracy of DBH estimations using backpack LiDAR resulted in a root mean square error (RMSE) of 2.7 cm and a coefficient of determination of 0.67. Conversely, the accuracy achieved with UAV LiDAR was 4.0 cm in RMSE and a 0.24 coefficient of determination. The heritability of DBH was found to be higher for backpack LiDAR than for UAV LiDAR and even exceeded that of hand measurements. Comparisons of the accuracy of timber volume selections based on the measured traits demonstrated comparable performances between the backpack and UAV LiDAR. Overall, these findings underscore the potential of using LiDAR remote sensing to quantitatively measure forest tree biomass and facilitate their genetic improvement of carbon-sequestration ability based on these measurements.

## 1. Introduction

In recent years, the importance of forests as carbon sinks has been increasingly recognized, particularly in the context of climate change mitigation and adaptation [1–4]. Plant biomass growth is primarily influenced by a combination of environmental and genetic factors. Quantitative genetics, a discipline that focuses on understanding the interplay between genetic and environmental factors in shaping the phenotypic traits of organisms, aims to quantify the extent to which trait variations are attributed to genetic inheritance. This framework of quantitative genetics serves as a powerful tool for dissecting the drivers of forest biomass growth, thereby enabling differentiation between the contributions of genetic and environmental factors. Moreover, it provides insights into the extent to which biomass growth can be influenced by human-mediated genetic interventions such as tree breeding.

In quantitative genetics, genetic relationships among individuals and their phenotypic values for a particular trait are collected and analyzed to ascertain the genetic contribution to trait variation. Traditionally, parent-progeny relationships and pedigree relationships over multiple generations have been used as genetic relationships among individuals. Recent developments in next-generation sequencing technology have enabled us to read a large number of DNA polymorphisms in many individuals and infer genetic relationships based on these polymorphisms. However, measuring phenotypic data is limited by the number of individuals that can be studied using conventional manual measurements, thereby making labor-saving measurements a challenge. Quantitative genetic analysis requires high efficiency and a certain level of accuracy in measuring a large number of individuals. The required accuracy level depends on the traits, population, and purpose of the analysis. Even if the measurement accuracy of an individual decreases, the ability to measure a large number of individuals can statistically compensate for the decrease in measurement accuracy of an individual. Against this background, remote sensing (RS) using unmanned aerial vehicles (UAV) and other high-throughput phenotyping methods is increasingly used for quantitative genetic analysis [5–10].

Light detection and ranging (LiDAR) technology, which has recently been increasingly used in combination with UAV RS, has been studied for application in forest management. In particular, the estimation of diameter at breast height (DBH) has been the subject of many studies because of its importance in forest management and the inefficiency of manual measurements [5, 6, 7]. To accurately determine the desired plant characteristics (e.g., DBH and tree height) from data collected by RS, developing data analysis methods is necessary, and many studies have been conducted to improve measurement accuracy [5, 11–17]. The results of these studies are important for the use of RS technology in plant phenotyping and quantitative genetics. However, few studies have examined the effectiveness and accuracy of RS for phenotypic measurements in quantitative genetic analysis of forest trees. Therefore, the objective of this study was to investigate the potential applications of LiDAR RS in quantitative genetic studies of forest trees. In particular, this study used two different LiDAR platforms—backpack and UAV—and compared their accuracies. The UAV-based LiDAR RS is highly efficient and covers a wide area. LiDAR measurement using a backpack is a relatively new technique that acquires data by walking on the forest floor using a sensor attached to the backpack. Although it is inferior to UAVs in terms of measurement efficiency, it has mobility that TLS does not have and is advantageous for acquiring data on forest structure, such as terrestrial laser scanning (TLS), under the canopy.

Larch, a deciduous conifer, constitutes approximately 30% of Japan’s forest area according to Forestry Agency statistics for 2021 (https://www.rinya.maff.go.jp/j/kikaku/toukei/attach/pdf/youran_mokuzi2023-3.pdf,/Japanese) and is one of the major plantation species in Japan [18]. Since the 1950s, larches have been selectively bred to promote strong and vigorous growth and the formation of straight trunks [18]. Notably, larches play a significant role in carbon sequestration, and Hirano et al. (2003) demonstrated that larch forests in Japan act as carbon sinks through negative net ecosystem CO2 exchange [19]. The deciduous characteristics of larches are particularly important for UAV LiDAR measurements because they facilitate trunk scanning without interference from the tree canopy during defoliation seasons. Consequently, larch is an ideal tree species for assessing the measurement accuracy of LiDAR across various platforms.

In this study, we aimed to develop methods for rapid and accurate measurement of larch biomass growth using LiDAR RS. We hypothesized that the accuracy of measurements would vary depending on the LiDAR platform, considering the sensor’s ranging accuracy. To test this hypothesis, we used a genetic model based on genome-wide polymorphism data. Specifically, we established methods for measuring tree location, diameter, and height using LiDAR RS for 602 60-year-old larch trees. These methods were applied using both backpack and UAV LiDAR RS platforms, and their capabilities to measure tree height and accuracy of diameter estimation were evaluated. Genome-wide marker data were collected from the target trees to determine their genetic relationships. Using these data, we estimated the heritability of traits and genetic correlations for different combinations of measurement methods and traits. Subsequently, we assessed the potential of LiDAR RS data for the efficient selection of timber volume. Through these investigations, we aimed to evaluate the potential of LiDAR RS measurements to assess forest tree biomass and contribute to genetic improvement efforts from a quantitative genetic perspective.

## 2. Materials and Methods

### 2.1. Study site

The 602 larch trees examined in this study were planted within the test site for hybrid larches at the Fuji Iyashinomori Woodland Study Center located in Yamanakako-mura, Minamitsuru-gun, Yamanashi, Japan. The region experiences average annual precipitation of 2,355 mm and an average annual temperature of 9.9°C, with a minimum temperature of −19.4°C, as recorded by the Amedas Yamanaka observatory. The soil in the area is classified as a gravelly volcanic immature soil, and the terrain is gently sloping.

The test site for hybrid larches was initially planted in 1955 with 1,806 individuals from 15 families. At present, 602 individuals remain as trees of approximately 60 years of age. The trees were arranged in a grid consisting of 24 rows and 76 columns spaced 2 m apart (Figure 1A). No thinning was conducted after the initial planting. Regular management operations removed the majority of understory vegetation from the forest floor.

**Fig. 1.**
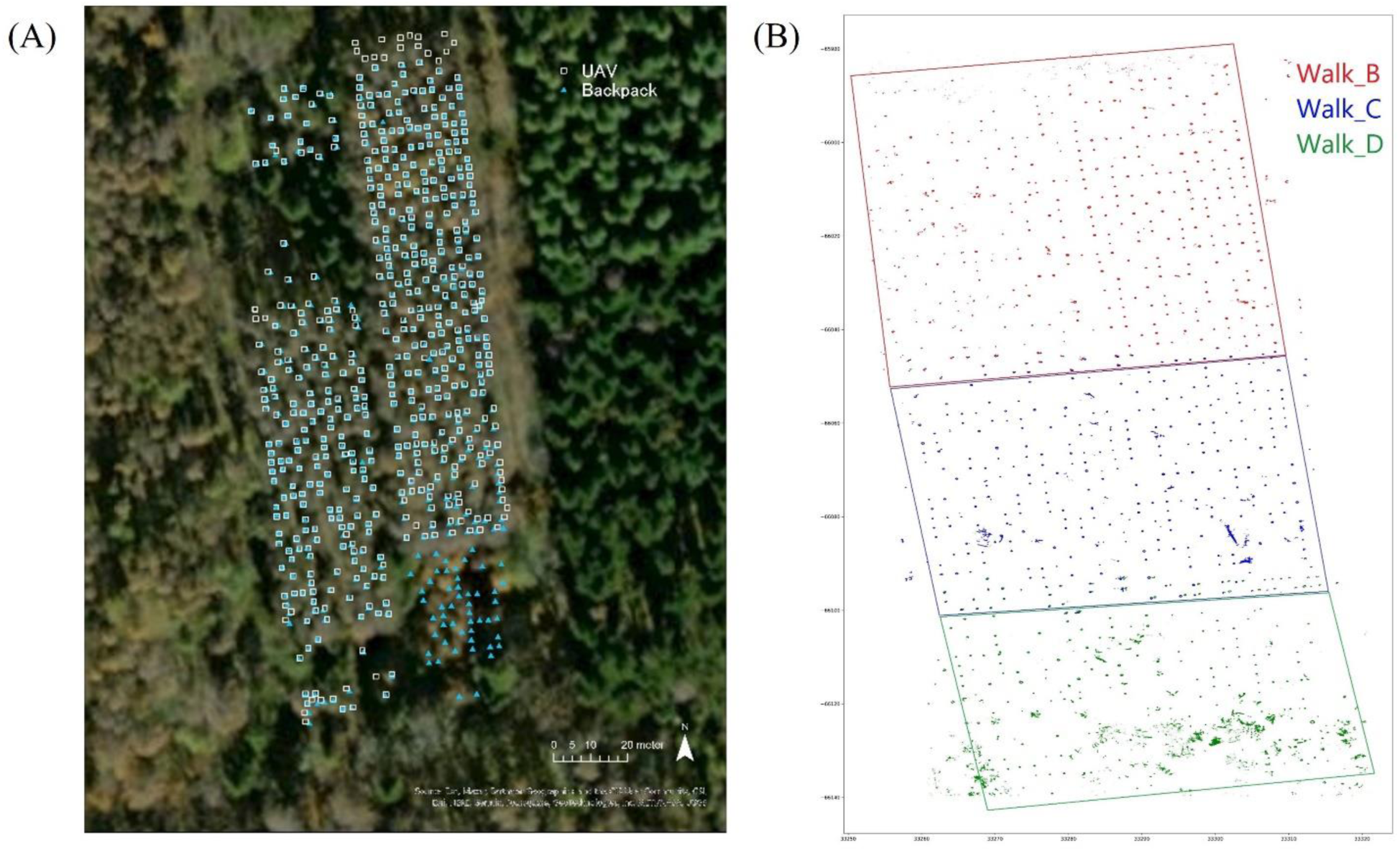
Overview of the study site. (A) Location map of each individual tree. Individual tree locations were estimated by analyzing point clouds obtained from backpack LiDAR (depicted as light blue triangles) and UAV LiDAR (depicted as white squares). (B) Trunk point overview of backpack LiDAR. The measurement area was divided into three sections in the north-south direction, from which three sets of point cloud data were acquired: Walk_B (depicted in red), Walk_C (depicted in blue), and Walk_D (depicted in green). Points at a height of 1.1–1.3 m above ground level were extracted from the acquired pint clouds and are represented here as “trunk points”. Abbreviations: LiDAR, light detection and ranging; UAV, unmanned aerial vehicle.

### 2.2. Hand-measured record of DBH

Manual measurement records of DBH were available for 205 individuals from seven families, measured using climbing plants (*Toxicodendron orientale* Greene and *Helianthus petiolaris* Siebold et Zucc.) in 2020. Among these, three were identified as full-sib families. The 2020 records were considered the true values to verify the DBH measurement accuracy.

### 2.3. Point cloud data acquisition

The point cloud data were acquired using two types of laser scanners: backpack LiDAR and UAV LiDAR. Table 1 presents the details of the sensors used. Both sets of point clouds were initially collected in the WGS 84 coordinate system and later transformed into Zone 8 of the Japanese plane rectangular coordinate system, where the unit is m, for analysis.

**Table 1.**
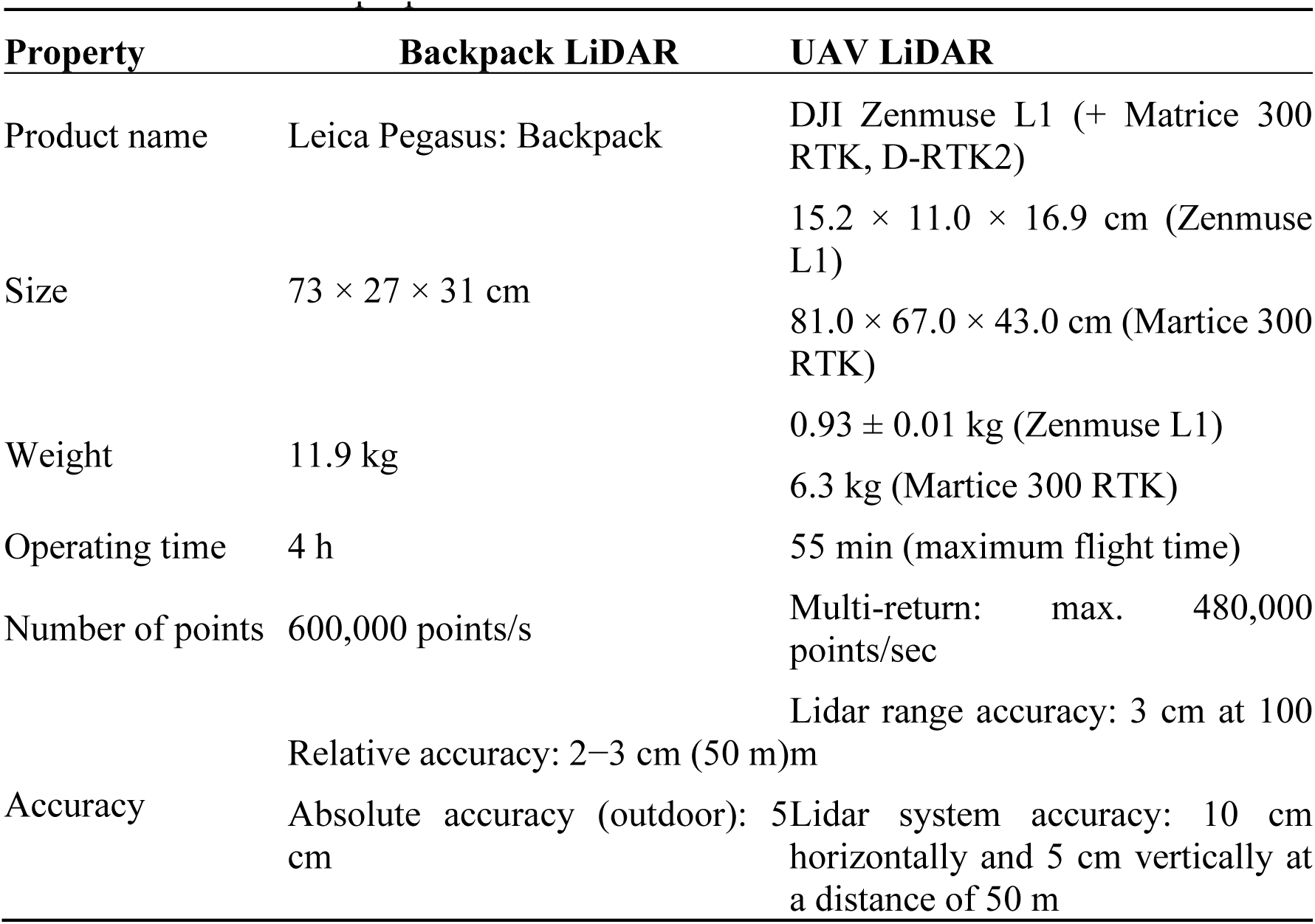
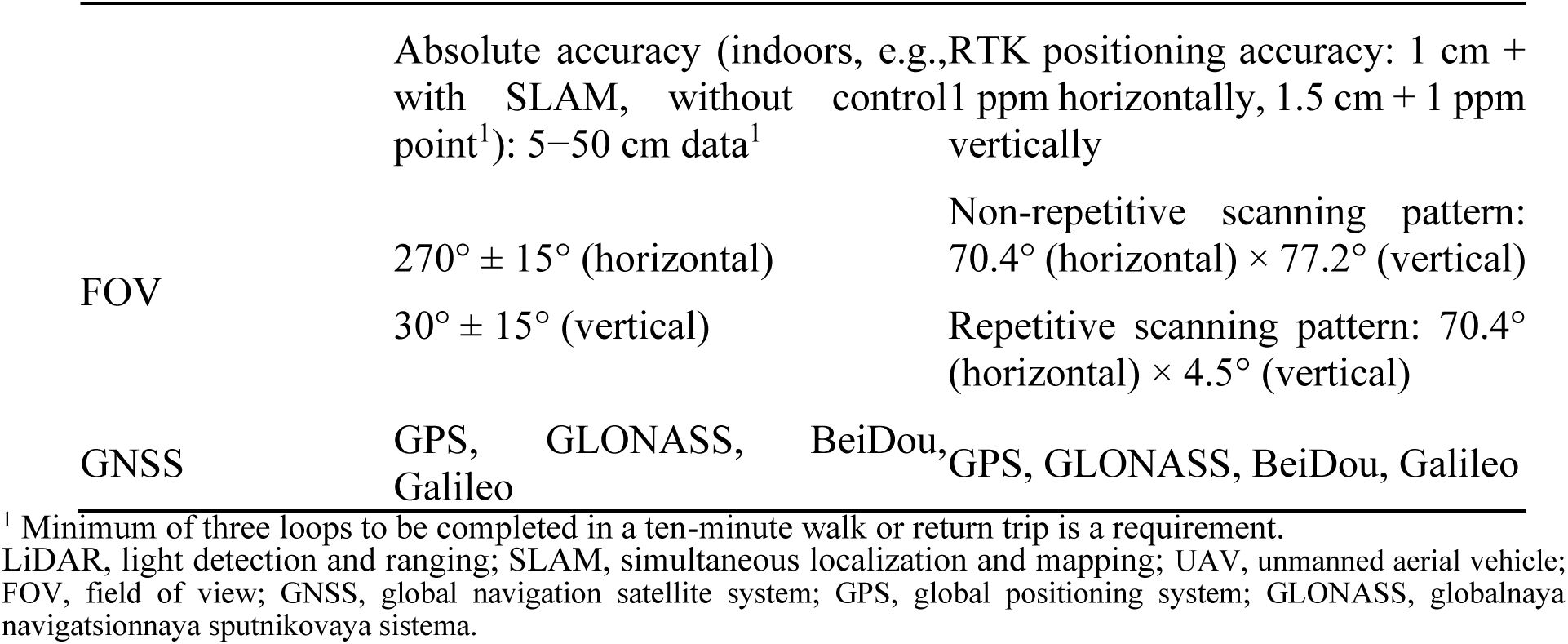
Laser scanner properties.

The backpack LiDAR used was the “Leica Pegasus: Backpack” manufactured by Leica Geosystems. This system uses two Velodyne VLP-16 scanners and is operated by carrying the scanners on a backpack while walking around the area to be measured. The relative accuracy of the system was reported to be 2–3 cm at a distance of 50 m with a pulse repetition rate of 60 kHz (30 kHz per scanner). Although the absolute accuracy when using the global navigation satellite system (GNSS) outdoors was 5 cm, measurements taken without control points based on simultaneous localization and mapping (SLAM) without the GNSS yielded accuracies ranging from 5 to 50 cm after 10 min of walking. In this study, data acquisition was performed using a combination of GNSS and SLAM techniques with seven control points. The test site was partitioned into three sections in the north-south direction, referred to as Walk_B, Walk_C, and Walk_D. Point cloud data were acquired for three sections on March 10, 2021. Specifically, rows 1–31 of the larch planting matrices were captured during Walk_B; rows 32–56 during Walk_C; and rows 57–76 during Walk_D (Figure 1B).

The UAV LiDAR was a Zenmuse L1 (DJI) mounted on a Matrice 300 RTK drone (DJI). This LiDAR system scans from the sky, with reported horizontal and vertical accuracies of 5 cm and 10 cm, respectively, at a distance of 50 m. Point cloud data were acquired from five directions, with gimbal angles of −90° and −45° (four directions), flight altitudes of 50 and 70 m, and returns set to dual and triple. The four patterns of data were combined after acquisition on March 2, 2022. The RTK positioning was performed using the DJI D-RTK2 system. The pulse repetition rate was 240 kHz for the dual return and 160 kHz for the triple return, with a 50% overlap between the flight paths.

### 2.4. Analysis flow of point cloud data

Point cloud data collected via backpack LiDAR in 2021 and UAV LiDAR in 2022 were analyzed separately to estimate the DBH and to compare the accuracy of the two LiDAR systems. The analysis flow is delineated in Figures S1–S4: Figure S1 illustrates the entire analysis flow, Figure S2 showcases the location estimation of individual trees, Figure S3 elaborates on the details of DBH estimation, and Figure S4 explains the tree height estimation.

First, noise was eliminated using a noise filter implemented in Cloud Compare [20], and the ground point clouds were classified using the Cloth Simulation Filter [21]. Subsequently, a digital terrain model (DTM) was generated using ordinary kriging. Point clouds situated at specific ground heights (1.1–1.3 m above ground for backpack LiDAR and 5–7 m above ground for UAV LiDAR) were extracted and identified as “trunk points” as depicted in Figure 1B. These trunk points were processed using Cloud Compare [20], from which the XY coordinates of the three corners of the planted square area were sampled (as outlined in Equation 1). Using these three points, a vector representing the movement of one row and one column (denoted as **R** and **C** in Equations 2 and 3, respectively) and the XY coordinates of the reference points of row 0 and column 0 (designated as p_0,0_ in Equation 4) were calculated. Based on these parameters, the approximate planting positions of the individual trees were estimated (designated as p_c,r_ in Equation 5).

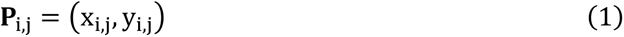

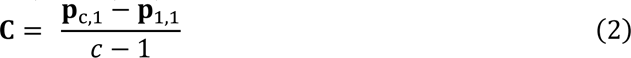

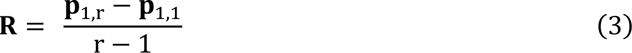

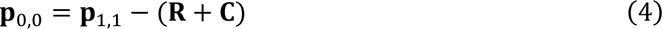

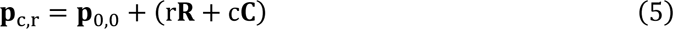

The planting location was then refined, starting with the initial approximate planting location. The center of gravity of the point cloud within a 0.6-m radius from the planting location was calculated, and this center of gravity was adopted as the new planting location. This process was iterated until the distance traveled in one step was less than 0.2 m or the distance from the initial value exceeded 1 m. Subsequently, manual position adjustments were made for 12 individuals using the backpack LiDAR and three individuals using the UAV LiDAR. The Z-coordinates of the ground at the individual planting positions were re-determined through kriging, using the XY coordinates of the individuals. For each individual, a point cloud (comprising trunk points) with a horizontal distance of less than 0.5 m from the planting location and at a certain height above the ground (1.1–1.3 m for the backpack LiDAR and 5–7 m for the UAV-type LiDAR) was extracted.

Subsequently, the XY coordinates of the tree trunk points were used to fit a circle and estimate the tree trunk radius (Figure 2). Specifically, we minimized the function v(b, **c**) described in Equation 5, where b represents the circle radius, and **c** = (c_**x**_, c_**y**_) denotes the center XY coordinates. Function v(b, **c**) comprises two components: v_1_(b, **c**) and v_2_(b, **c**). The function v_1_(b, **c**) quantifies the percentage of points within a distance b ± w from the center of the fitted circle, where w is a ranging error adjustment parameter (set as 0.035). The function v_2_(b, **c**) evaluates the directional variation of the points from the center of the circle, calculated as one minus the average magnitude of the position vectors from the center of the fitted circle for points falling within b ± w. To perform the minimization, we used the fminsearch function [22] in MATLAB [23], which facilitates the estimation of the minimum of an unconstrained multivariable function without derivatives. The initial values of **c** and b were set as the average of the center of gravity of the trunk point and its distance from it for each individual.

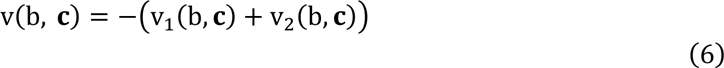

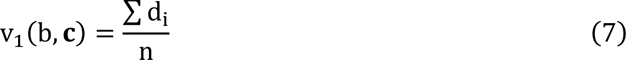

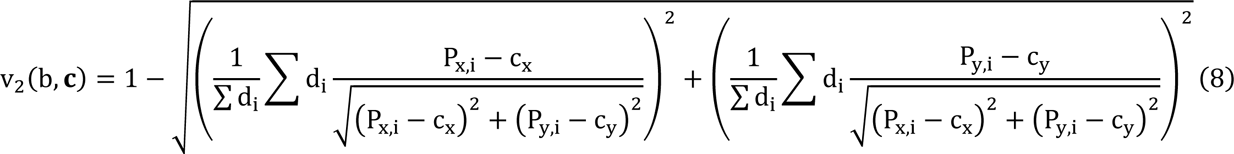

**Fig 2.**
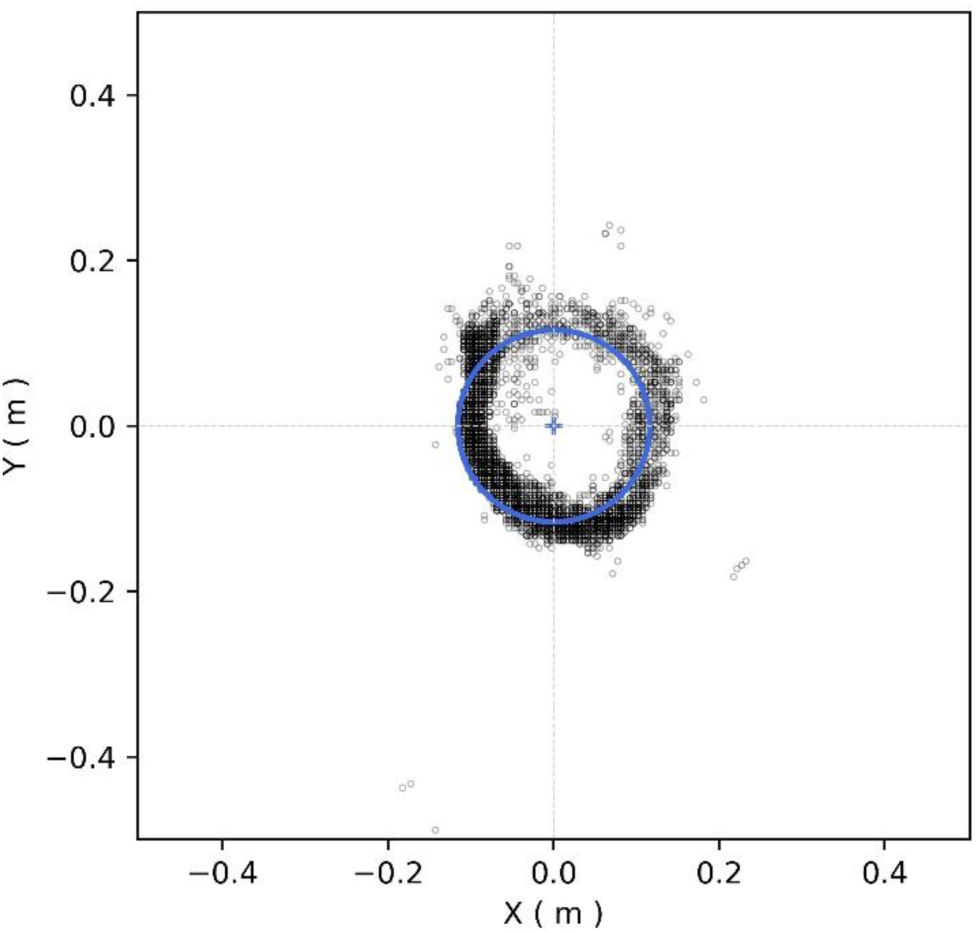
Example of circle fitting. The point cloud extracted from the backpack light detection and ranging (LiDAR) data for individual 247, situated at a height of 1.1–1.3 m above ground level, is represented by the black circle. The blue circle depicts the circle fitted to the data, whereas the blue cross denotes its center. Coordinates are extracted from the center of the fitted circle.

In Equations 6 and 7, P_x,i_and P_y,i_ represent the X- and Y-coordinates of the trunk point, respectively, n denotes the number of trunk points, and d_i_ is defined as follows: d_i_ takes the value of 1 when point P_i_ satisfies the condition 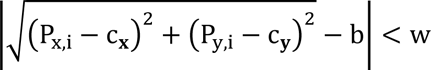, and 0 when it does not.

The measurement accuracy was assessed through linear regression, using the trunk diameter estimated using the method described above as the response variable and hand measurements as the explanatory variable (Equations 9 and 10). The root mean square error (RMSE) and coefficient of determination were calculated. For backpack LiDAR, individuals with an estimated diameter greater than 40 cm were identified as outliers and excluded from the analysis.

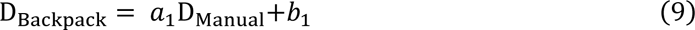

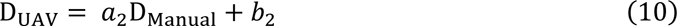

### 2.5. Genotyping of genome-wide high-density markers

Genome-wide marker genotypes were acquired through the RAD-Seq analysis of 333 trees using DNA samples. Genomic DNA was extracted from young leaf tissues using a DNeasy Plant Mini Kit (QIAGEN, Hilden, Germany) following the manufacturer’s protocol. Base variants were identified using dd-RAD-Seq reads. Library construction was performed as described by Shirasawa et al. [24], and dd-RAD-Seq sequences were obtained using the DNBSEQ-G400RS platform (MGI Tech Co., Ltd., Shenzhen, China). The reads were aligned to the published Japanese larch (*Larix kaempferi*) genome [25] using Bowtie 2 [26]. A variant call was conducted using bcftools 0.1.19 mpileup in SAMtools [27] and variant quality filtering was performed using vcftools [28]. Imputation of missing entries in the marker genotype data was performed using Beagle 5.4 [29].

### 2.6. Estimation of heritability

Genomic heritability was estimated using the genome-wide marker data for each DBH measured using the three methods: the DBH from hand-measured records in 2020 and the estimated DBH from backpack and UAV LiDAR. Data from 43 individuals with genome-wide markers and complete phenotypic data, which indicated that there were no missing entries across the four traits (three DBH measurements and tree height), were used for the estimation. In addition, the heritability of tree height was estimated from the UAV LiDAR point cloud using the same individuals.

To estimate genomic heritability, markers with a frequency of three or fewer individuals, corresponding to a minor allele frequency of less than 0.02% were initially excluded from the genome-wide marker genotype data. Subsequently, the marker genotype scores (0, 1, 2) were standardized to have a mean of zero and a variance of one and were arranged into a matrix format as **X**. Rows represented individuals, and columns represented markers. The genomic relationship matrix, denoted by **G**, was derived from **X** according to Equation 11.

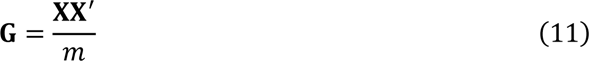

In Equation 11, *m* represents the number of markers. The R package BGLR [30] was used to estimate the parameters of a Bayesian-generalized linear regression model for each of the four traits (DBH measured using the three methods and tree height) serving as response variables. Matrix **G** was used as the covariance structure of the multi-normal distribution to which genetic values, specifically breeding values, adhered (**g** in Equation 12).

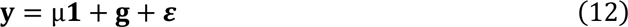

In Equation 12, **y** represents the vector of the response variable (DBH or tree height), μ stands for the overall mean, **1** is a vector of elements equal to 1 with dimensions matching the number of individuals (n), **g** signifies the vector of breeding values accounted for by the genomic relationships, assuming a multivariate normal distribution **g**∼N(0, **G***σ*_*g*_), and *ε* represents the vector of residuals, assuming a multivariate normal distribution *ε*∼N(0, **I***σ*_*e*_). **I** denotes the n × n-dimensional identity matrix.

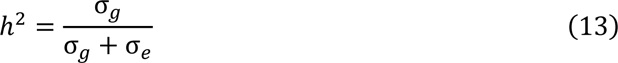

The ratio of genetic variance to total variance (genetic variance σ_*g*_ + environmental variance σ_*e*_) was estimated as the genomic heritability, as shown in Equation 13.

### 2.7. Genetic correlation and timber volume selection accuracy

To assess the accuracy of timber volume selection based on remotely sensed traits, we estimated the genetic correlations between these traits, including timber volume. The calculations followed the methods outlined by Xu [31], with reference to previous studies [32–34].

First, a multi-trait model (Equation 14) was applied, where the phenotypes comprised logarithms of DBH measured by hand, backpack LiDAR, and UAV-type LiDAR, as well as tree height values. This model aimed to estimate heritability, genetic variance, genetic correlation, and genetic variance-covariance matrices. Notably, logarithmic values of DBH and tree height were used for subsequent calculations.

In Equation 14, **y** represents the vector with a length equal to the number of individuals multiplied by the number of traits (four), **µ** denotes the vector whose elements correspond with the overall mean, *ε* represents the vector of residuals, and **g** stands for the vector of genetic effects, taking into account the covariance between traits and the genomic relationships, i.e. **g**∼N(0, ∑×**G***σ*_g_^2^), where ∑ is the among-trait genetic covariance matrix to be estimated and × is the kronecker product.

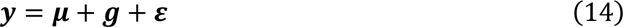

We assessed the accuracy of timber volume selection based on DBH (*D*) and tree height (*H*). Assuming that the main trunk of a larch tree is conical, timber volume (*V*) can be expressed as *V* = *CD*^2^*H*, where *C* is a constant. We modeled the logarithmic expression of timber volume as *S* = log *V* = 2 log *D* + log *H* + *C* , using tree height derived from the point cloud and DBH value. Here, we assumed that *S* could be calculated using the hand-measured DBH and that the estimated tree height was the true value.

Subsequently, we derived an optimized equation for inferring *S* (referred to as selection index I; see Equation 14) indirectly using a linear combination of measurements in the three patterns to evaluate the accuracy of timber volume selection. Pattern 1 involved hand-measured DBH and tree height, Pattern 2 involved DBH measured with backpack LiDAR and tree height, and Pattern 3 involved DBH measured with UAV LiDAR and tree height.

X represented a measurement pattern, P the covariance matrix of the estimated values of the three traits, and K the genetic covariance matrix. Phenotypes were calculated after normalization to a mean of zero and a variance of one. The optimal weight b can be expressed as shown in Equation 15, where w is the weight of the target trait, with two assigned to DBH hand measurements and one to tree height measurements, according to the model equation for timber volume.

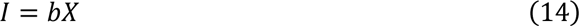

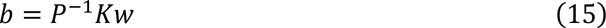

The value of b represents the weight given to the measurements in each pattern to estimate the genetic ability of timber volume. Subsequently, the correlation coefficient between the selection index I and target trait S was calculated (Equation 16–18). The correlation coefficient r represents the accuracy of estimating the genetic ability of timber volume, i.e., the accuracy of timber volume selection based on the estimation. These values were calculated for the three patterns and compared.

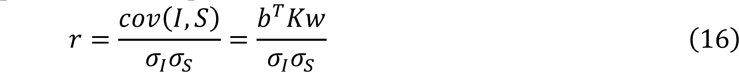

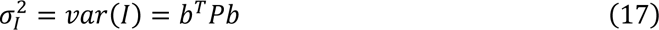

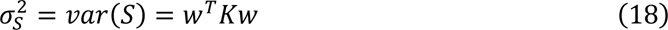

## 3. Results

### 3.1. Tree location estimations

As a result of the planting location estimation of individual trees, 578 of 602 trees could be estimated with backpack LiDAR, whereas 546 of 602 trees could be estimated with UAV LiDAR (Figure 1A). Among them, 524 trees were estimated using both methods. For these 524 trees, the mean, median, and standard deviation of the distances between the planting locations estimated using the backpack and UAV LiDAR were 0.32 m, 0.26 m, and 0.25 m, respectively. As depicted in Figure 1A, the backpack LiDAR failed to estimate the location in the northeastern region, most likely owing to the lack of ground points in that region. Conversely, the UAV LiDAR failed in the southeastern area, possibly because of the low tree height (2–3 m).

### 3.2. Accuracy of DBH estimation

The accuracy of the DBH estimation obtained using both backpack and UAV RS was validated against hand measurements (Figure 3). The estimation accuracy yielded RMSE of 2.7 cm and 4.0 cm, with coefficient of determination values of 0.67 and 0.24 for the backpack and UAV RS, respectively. Notably, the backpack RS exhibited superior accuracy compared with that of the UAV RS. Mean, median, and standard deviation of the errors were recorded as −2.6 × 10^−15^ cm, 0.01 cm, and 2.7 cm for the backpack RS and 3.2 × 10^−15^ cm, −0.9 cm, and 4.0 cm for the UAV RS, respectively. Figure 4 illustrates histograms of the errors and their kernel densities. Major error sources included climbing plants entwined around trunks as well as understory vegetation and shrubs proximal to the trunks.

**Fig 3.**
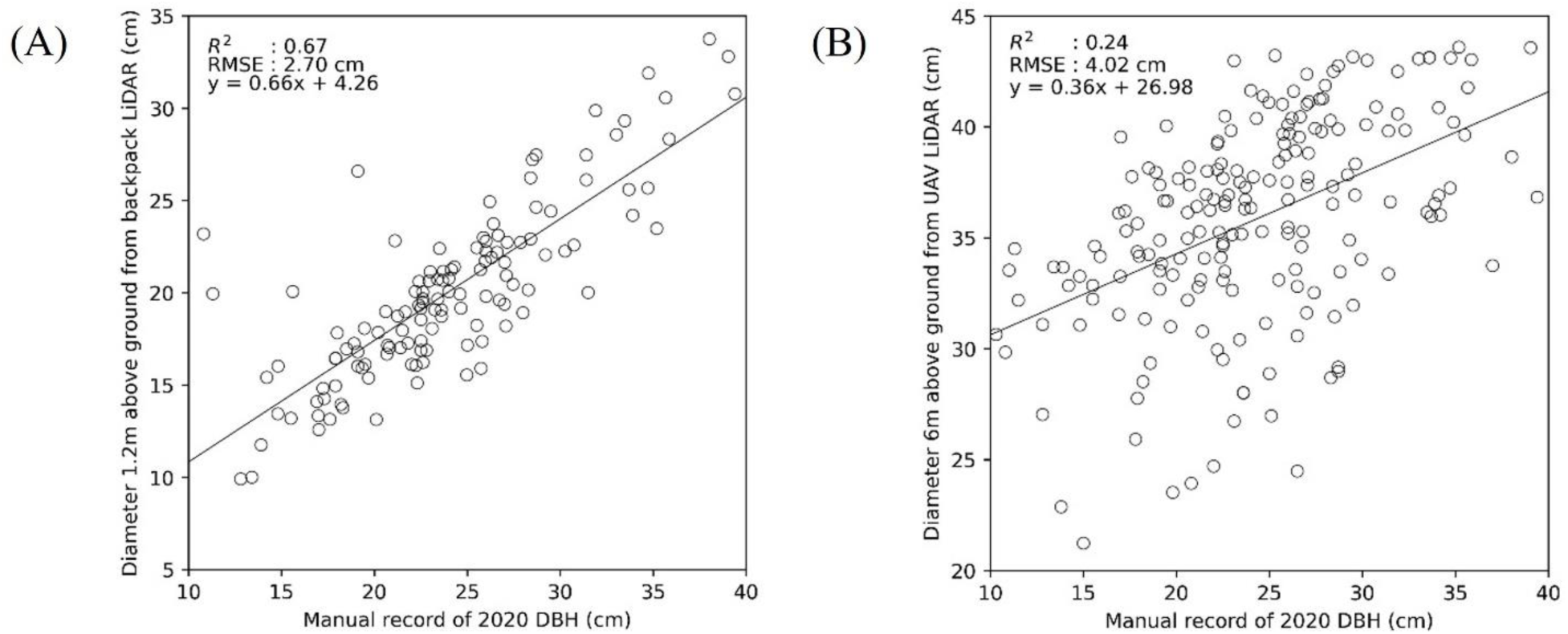
Comparison of estimated diameter with reference values. (A) Diameter estimated using backpack LiDAR and hand measurements. The horizontal axis represents the hand-measured record for 2020, whereas the vertical axis depicts the estimated diameter using backpack LiDAR. (B) Comparison of DBH estimated using UAV LiDAR and hand measurements. The horizontal axis represents the hand-measured record for 2020, and the vertical axis is the estimated diameter using UAV LiDAR. Abbreviations: LiDAR, light detection and ranging; UAV, unmanned aerial vehicle; DBH, diameter at breast height.

**Fig 4.**
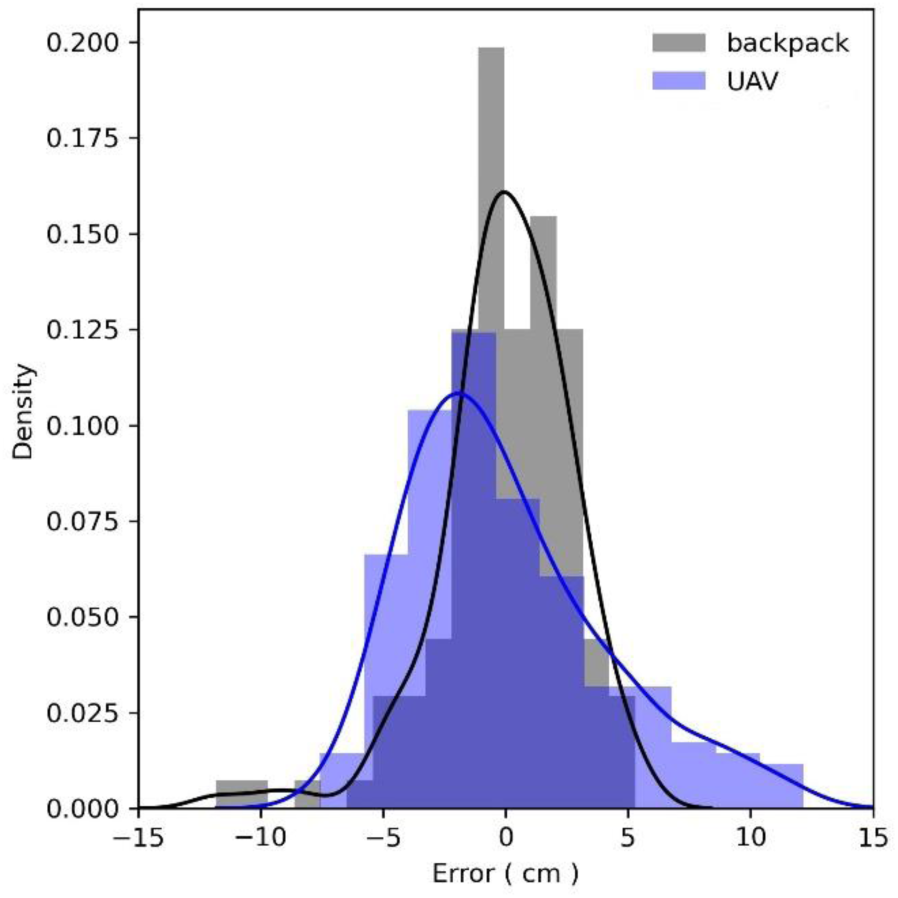
Histograms and kernel densities of DBH estimation. The errors are depicted for both the backpack LiDAR (gray) and UAV LiDAR (blue). Abbreviations: LiDAR, light detection and ranging; UAV, unmanned aerial vehicle; DBH, diameter at breast height.

### 3.3. Heritability of DBH and tree height

The heritability of the manually measured DBH was found to be 0.60. In comparison, the heritability of DBH estimates obtained through backpack LiDAR and UAV LiDAR were 0.62 and 0.53, respectively. The heritability of DBH with backpack LiDAR surpassed that of manually measured DBH. Additionally, the heritability of tree height estimated using UAV LiDAR was 0.52.

### 3.4. Genetic correlation among traits and accuracy of timber volume selection

Genetic correlation between manually measured DBH and DBH estimated with backpack and UAV LiDAR wad 0.82 and 0.57, respectively. The genetic correlation of tree height was 0.74 for manually measured DBH, whereas it was 0.66 and 0.55 for DBH estimated using backpack and UAV LiDAR, respectively.

To assess the accuracy of the genetic ability estimation for timber volume, the correlation between the selection index and target trait V was calculated for three combinations of tree height and DBH measurements using different methods.

Correlation coefficients (r) between selection index I and target trait V were 0.75 for pattern A using hand measurements, 0.78 for pattern B using backpack LiDAR, and 0.71 for pattern C using UAV LiDAR for DBH measurements. Notably, pattern B, which used backpack LiDAR, exhibited the highest correlation.

The weights (b) in selection index I were determined to be 3.50 and 3.55 for DBH and tree height, respectively, for pattern A, 4.05 and 4.12 for pattern B, and 3.49 and 4.01 for pattern C. Patterns B and C, which were based on LiDAR-based DBH measurements, indicate that selection should assign a stronger weight to DBH than to tree height.

## 4. Discussion

### 4.1. Comparison of accuracy with previous studies

This study evaluated two different platforms for measuring tree biomass-related traits, namely, DBH and tree height. Previous studies on DBH measurement using LiDAR can be categorized as those using airborne laser scanning (ALS), primarily with aircraft or UAVs as platforms, TLS or Mobile Laser Scanning (MLS). The backpack LiDAR used in this study falls under MLS, whereas the UAV LiDAR belongs to ALS.

Among studies using ALS to measure DBH, Brede et al. [11] reported an accuracy of 4.24 cm for RMSE (using TLS estimates as the reference value), Yao et al. [12] documented 4.15 cm for coniferous trees and 5.07 cm for deciduous trees, Popescu et al. [13] found 4.9 cm, and Dalla Corte et al. [5] achieved 3.46 cm. Previous studies using TLS include Huabing et al. [15] with 3.40 cm, Monika et al. [16] with 9.17 cm, Sanzhang et al. [17] with 0.70 cm, and Liu et al. [14] with 3.17 cm for natural forest and 1.97 cm for urban forests. Additionally, Zhang et al. [35] reported an accuracy of 1.48 cm using backpack LiDAR. Although the accuracy of DBH estimation may vary depending on factors, such as tree species, individual characteristics, forest conditions, LiDAR accuracy, and algorithms used, the results of 2.3 cm for the backpack LiDAR and 4.0 cm for the UAV LiDAR in this study compare favorably with findings from previous studies.

### 4.2. Required accuracy level

Several factors must be considered when determining the requisite accuracy of the DBH measurements. First, assessing the ranging accuracy of LiDAR systems is imperative. Considering the nominal accuracies of the backpack LiDAR and UAV LiDAR used in this study, which were 2 – 3 cm and 5– 10 cm, respectively, the DBH measurement accuracies of 2.7 cm and 4.0 cm, respectively, are justifiable. Second, although manually measured DBH is typically rounded to the nearest centimeter, it is crucial to acknowledge that this does not imply a precision level of 1 cm. Variations in the orientation, height, and tilt of the caliper against the tree trunk can introduce significant error margins in manual measurements. Additionally, for accurate assessment of biomass accumulation in forests, simultaneous measurements of tree height and DBH are essential, as underscored in this study. Although the tree height measurement accuracy was not evaluated in this study owing to the absence of true values, future accuracy verification will be indispensable for precise biomass measurement.

Furthermore, in terms of forest tree breeding, heritability has emerged as a pivotal criterion for evaluating accuracy. A measurement method that exhibits high heritability, indicating a significant proportion of genetic variation among the observed variations, is required to enhance the efficiency of genetic improvement. Given that genomic heritability of DBH was estimated at 0.60 from hand measurements, 0.62 from backpack LiDAR, and 0.52 from UAV LiDAR, it became evident that backpack LiDAR provided a superior estimation of heritability compared with that of the UAV LiDAR. This disparity can be attributed to the difference in the DBH measurement accuracy, with 2.7 cm in backpack LiDAR and 4.0 cm in the UAV LiDAR.

### 4.3. Potential accuracy of timber volume selection

This study investigated the feasibility of accurately selecting timber volume as an additional criterion for evaluating the precision of LiDAR-based measurements of DBH and tree height.

Timber volume measurement is pivotal in forest management, closely linked to estimating forest carbon sequestration, and is a key target for natural forest trees. In this study, timber volume was defined as 2log(D) + log(T) + C, based on tree height (H) and hand-measured DBH (D), serving as the improvement target V. Subsequently, the optimal selection index for V was derived from the linear combination of tree height and DBH measured by the two types of LiDAR and manually, and the correlation between the selection index and genetic variation of V was calculated. Three measurement patterns were assumed for the selection index calculation: Pattern A used hand-measured DBH as D, Pattern B used DBH measured with backpack LiDAR as D, and Pattern C used DBH measured with UAV LiDAR as D. Finally, we assessed how the accuracy of the genetic variation estimation in the selection index varied across these three patterns, accounting for the genetic correlations among the traits.

The results revealed that the index of selection accuracy, r (correlation between the selection index and genetic variation of V), was 0.75 for hand measurements, 0.78 for backpack LiDAR, and 0.71 for UAV LiDAR. Compared with that of the hand measurements, the accuracy was the highest when backpack LiDAR was used and was slightly less accurate when UAV LiDAR was used. However, there was a marginal difference in the accuracy among the three methods. These findings indicate that both backpack LiDAR and UAV LiDAR exhibit similar performance in timber volume selection, suggesting a comparable selection accuracy between the two methods. Moreover, using UAV LiDAR for timber volume selection is highly practical, as it allows for the simultaneous measurement of tree height and DBH, streamlining the process.

When the target trait is derived from individually measured traits, selection accuracy cannot be solely determined by the accuracy of the individual measurements. Optimal measurement methods can be explored by conducting the calculations presented in this study. In particular, when a target trait can be evaluated based on traits that have not been conventionally used, such as DBH measured by LiDAR and UAV, performing evaluations akin to those in this study becomes crucial.

### 4.4. Differences between platforms and conditions for acquiring point cloud data

The acquisition and analysis of point clouds using two different platforms, a backpack and UAV, revealed notable differences between the platforms. These variances encompass the concentration of point clouds in specific regions, resulting in discrepancies in the measurable features, timeframes, and tree species.

The backpack platform acquires point clouds using a laser beam emitted approximately 2 m above the ground. Consequently, the point clouds were primarily concentrated on the ground and tree trunks. This setup renders it unsuitable for tree height estimation but is well suited for estimating DTMs and DBH. By contrast, the UAV platform emits lasers from above the area to be measured, resulting in point clouds concentrated within the tree canopy. Consequently, the point clouds of the trunk and ground diminish as the lasers are obstructed by the canopy. This configuration ensures highly accurate tree height measurements but results in less accurate DBH measurements. Brede et al. [11] also noted limitations in the detection of the top of the canopy using TLS. In the era of aircraft ALS, it was considered the most accurate method for measuring tree height among the previous methods; however, the underestimation of tree height remained (owing to laser pulses missing the top of the tree). A UAV with a dramatically increased density of points overcame this problem and enabled a more accurate tree height measurement.

Considering these characteristics, certain conditions for point cloud acquisition of diameter estimation have become apparent. In the case of deciduous coniferous trees, such as larches, UAV LiDAR measurements are recommended during the defoliation period. Conversely, in evergreen trees, the use of backpack LiDAR is preferable, at least based on the point cloud density and positioning accuracy observed in this study. Thus, the selection of a platform should be tailored to the specific features to be measured, resulting in optimal results.

### 4.5. Significance and comparison of location estimation

The ability to precisely measure the positions of individual trees using LiDAR represents a significant advantage, particularly in forested areas where GNSS reception is challenging under dense canopy cover. Accurately determining planting locations using LiDAR offers valuable insights, especially when integrated with other datasets such as passive remote sensing data [36]. In cases where trees are planted in a regular pattern, as in our study, we recommend using the method proposed herein for the semiautomatic estimation of the planting location of each individual tree.

Our study found that the estimated positions obtained from both the backpack and the UAV LiDAR exhibited good agreement, with an average difference of 0.3 m. However, the primary factor contributing to the positional discrepancies is the positioning accuracy of the backpack LiDAR. As illustrated in Figure 1B, there are instances in the point cloud data acquired from Walk_B and Walk_C of the backpack LiDAR where the trunk positions do not align. This discrepancy underscores the limitation of the positioning accuracy of backpack LiDAR without a GNSS. Pierzchała et al. [37] noted that improved SLAM techniques can enhance the accuracy of tree location estimation, suggesting promising avenues for refining positioning techniques such as SLAM.

In conclusion, our study explored the application of backpack LiDAR and UAV LiDAR for DBH measurement and genomic heritability estimation. Moreover, their accuracy was compared with that of hand measurements. We found that backpack LiDAR exhibited high accuracy, with an RMSE of 2.7 cm compared with 4.0 cm for the UAV Li-DAR. Furthermore, the heritability of DBH surpassed that of hand-measured DBH. We also assessed the accuracy of timber volume selection, revealing comparable performance between backpack LiDAR and UAV LiDAR.

These findings underscore the potential of LiDAR remote sensing for biomass growth measurements and genetic enhancement of the carbon sequestration ability of forest trees. Additionally, our analysis enabled the automatic estimation of tree locations for over 90% of the trees, leveraging coordinates from point cloud data and plantation maps. To fully evaluate the genetic potential of LiDAR remote sensing in forest management and tree breeding, further studies should investigate how various conditions, such as the presence of climbing plants and understory vegetation, planting density, tree height, and degree or absence of defoliation, affect measurement accuracy.

## Acknowledgments

The backpack LiDAR data used in this study were kindly provided by Leica Geosystems and we express our sincerest gratitude to Leica Geosystems. We would also like to thank the Fuji Iyashinomori Woodland Study Center and technical specialists Norio Nishiyama and Ryoko Tsuji for their cooperation in obtaining the data for the test site for hybrid larches, which is under the jurisdiction of the center. We are deeply thankful to Prof. Seiji Ishibashi for providing us with hand-measured DBH data. We also thank Akiko Watanabe, Yoshie Kishida, Hisano Tsuruoka, Keishi Ozawa, Taeko Shibazaki, and Yukio Matsumoto of Kazusa DNA Research Institute for their technical assistance.

## Author contributions

Conceptualization, H.S. and H.I.; methodology, H.S. and H.I.; software, H.S. and SI; validation, H.S.; formal analysis, H.S.; investigation, H.S., N.M., W.G., and H.I.; resources, K.K. and U.Y.; data curation, H.S. and N.M.; writing—original draft preparation, H.S.; writing—review and editing, H.I.; visualization, H.S.; supervision, H.I.; project administration, H.I.; and funding acquisition, H.I. All authors have read and agreed to the published version of the manuscript.

## Data Availability

Data will be made available on request.

## Supplementary Materials

### Supplementary figures

The flowchart of point cloud data analysis is presented below as Figures S1–S4.

**Fig S1.**
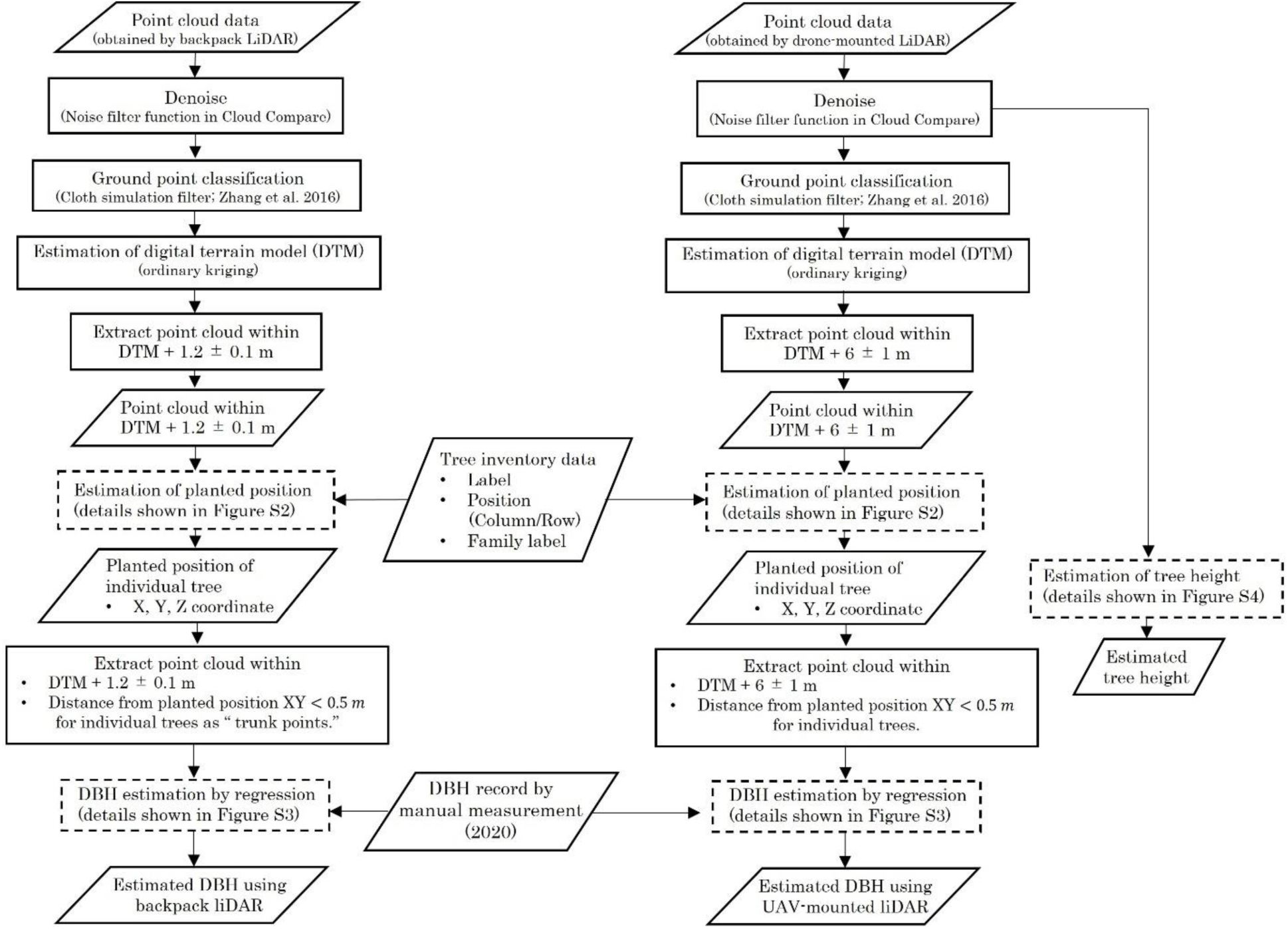
Analysis flow of point cloud data. The left side shows the procedure for estimating planting location and diameter at breast height (DBH) using point clouds acquired by backpack light detection and ranging (LiDAR) and by unmanned aerial vehicle (UAV) LiDAR on the right side.

**Fig S2.**
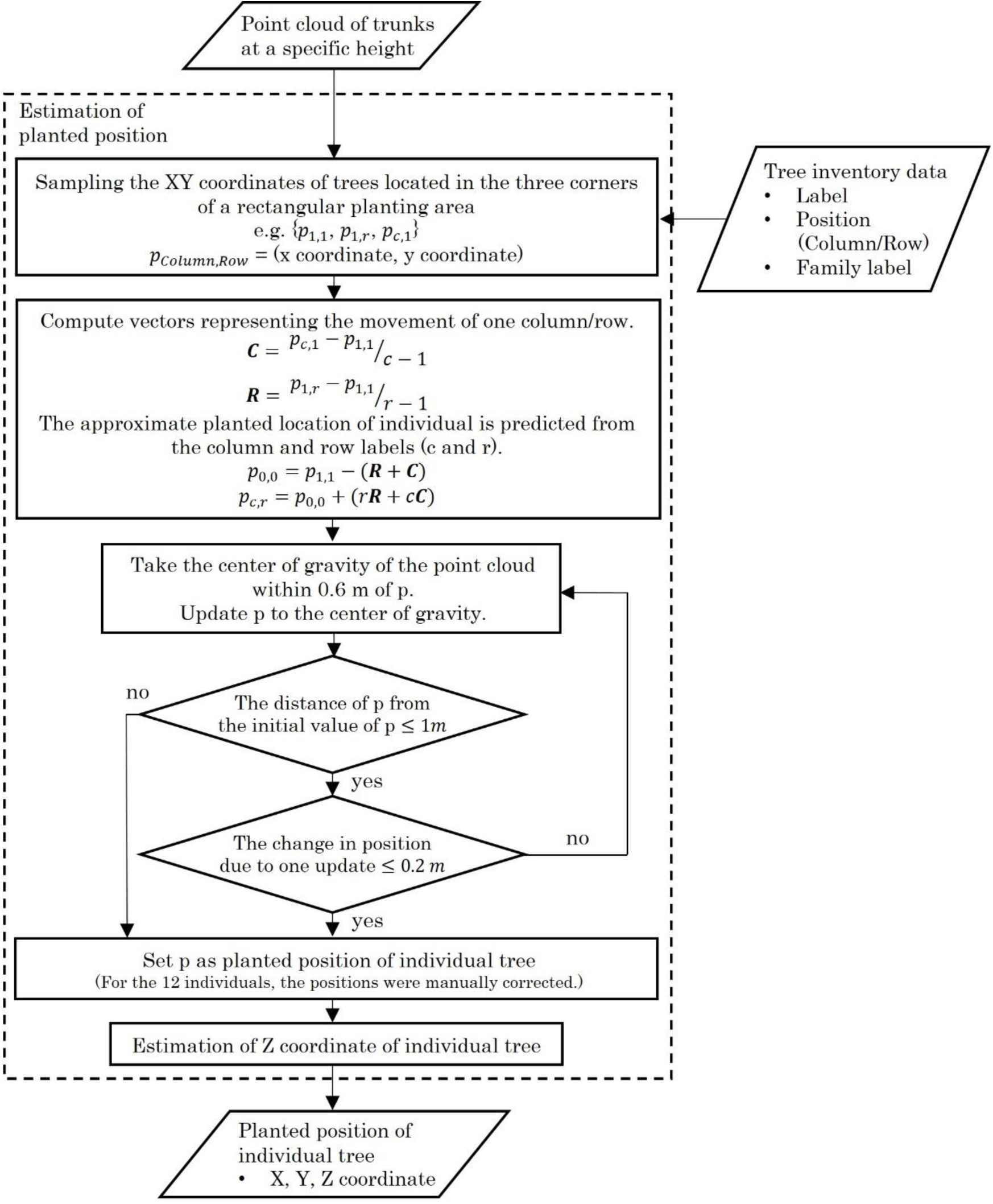
Individual location estimation in the point cloud data analysis flow.

**Fig S3.**
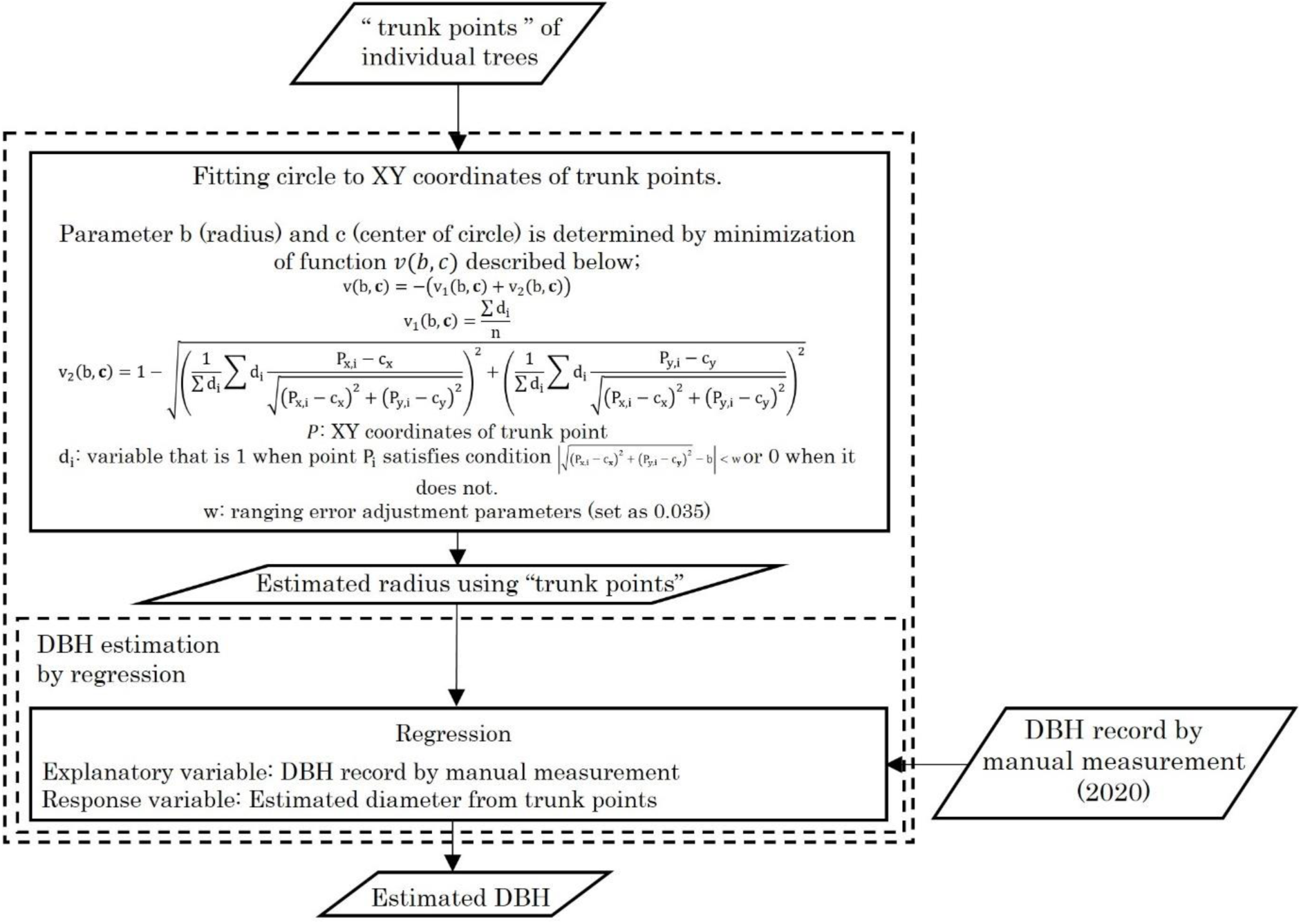
Diameter at breast height (DBH) estimation in the analysis flow of point cloud data.

**Fig S4.**
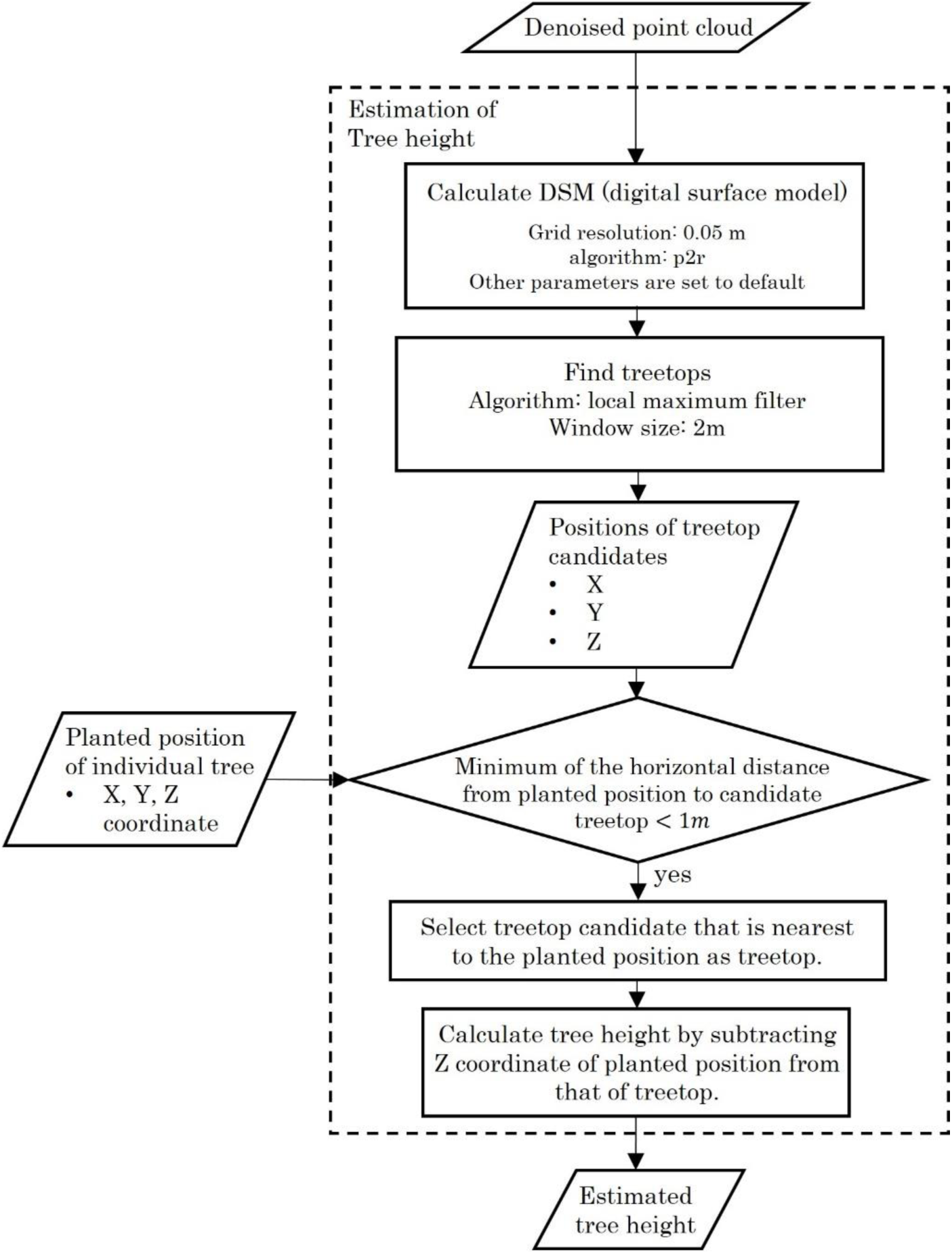
Tree height estimation in the analysis flow of point cloud data.

