## Supplementary figures and images for "Quantifying Forest Biomass and Genetic Contribution using Light Detection and Ranging"

### Supplemental figure 1

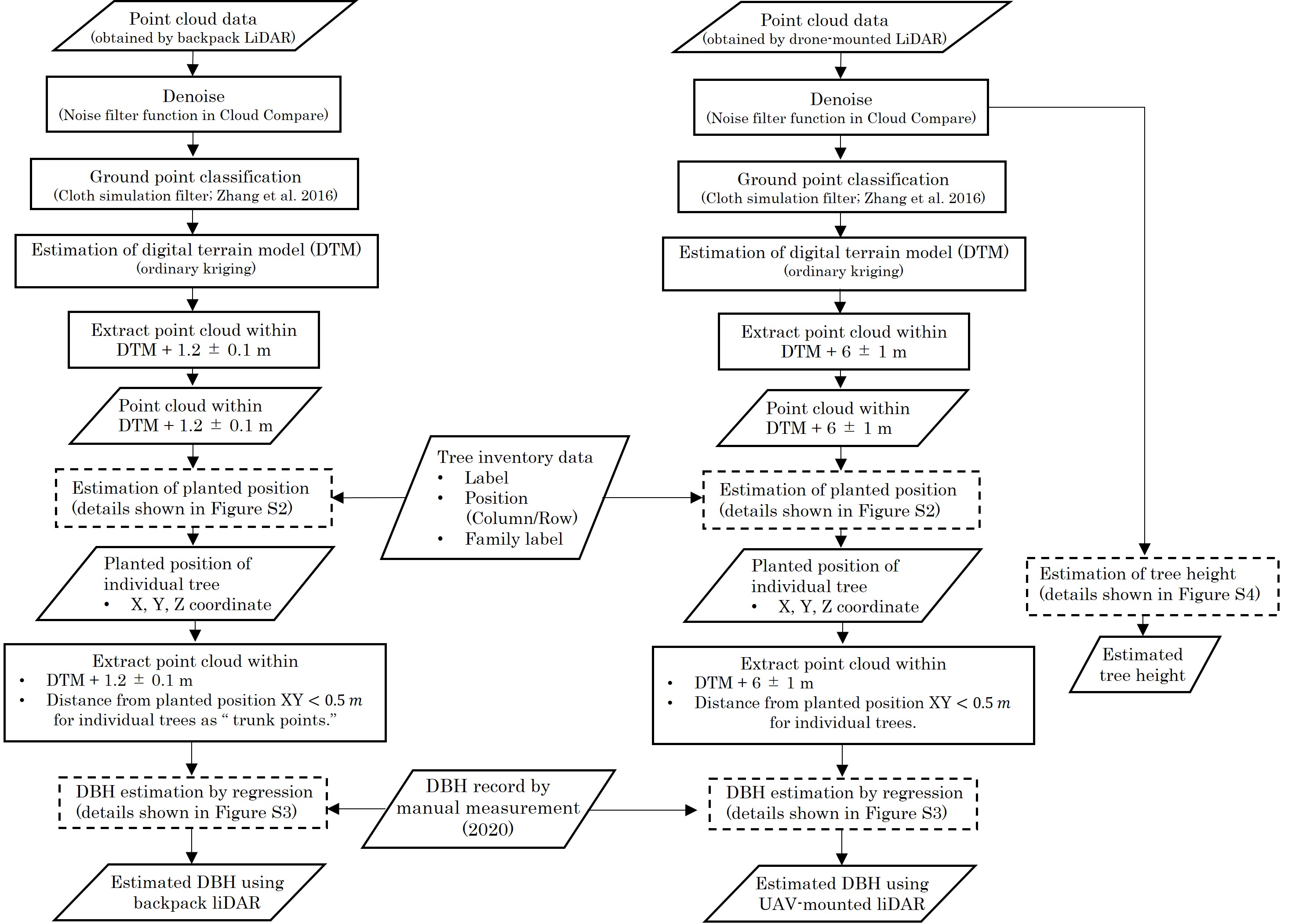

### Supplemental figure 2

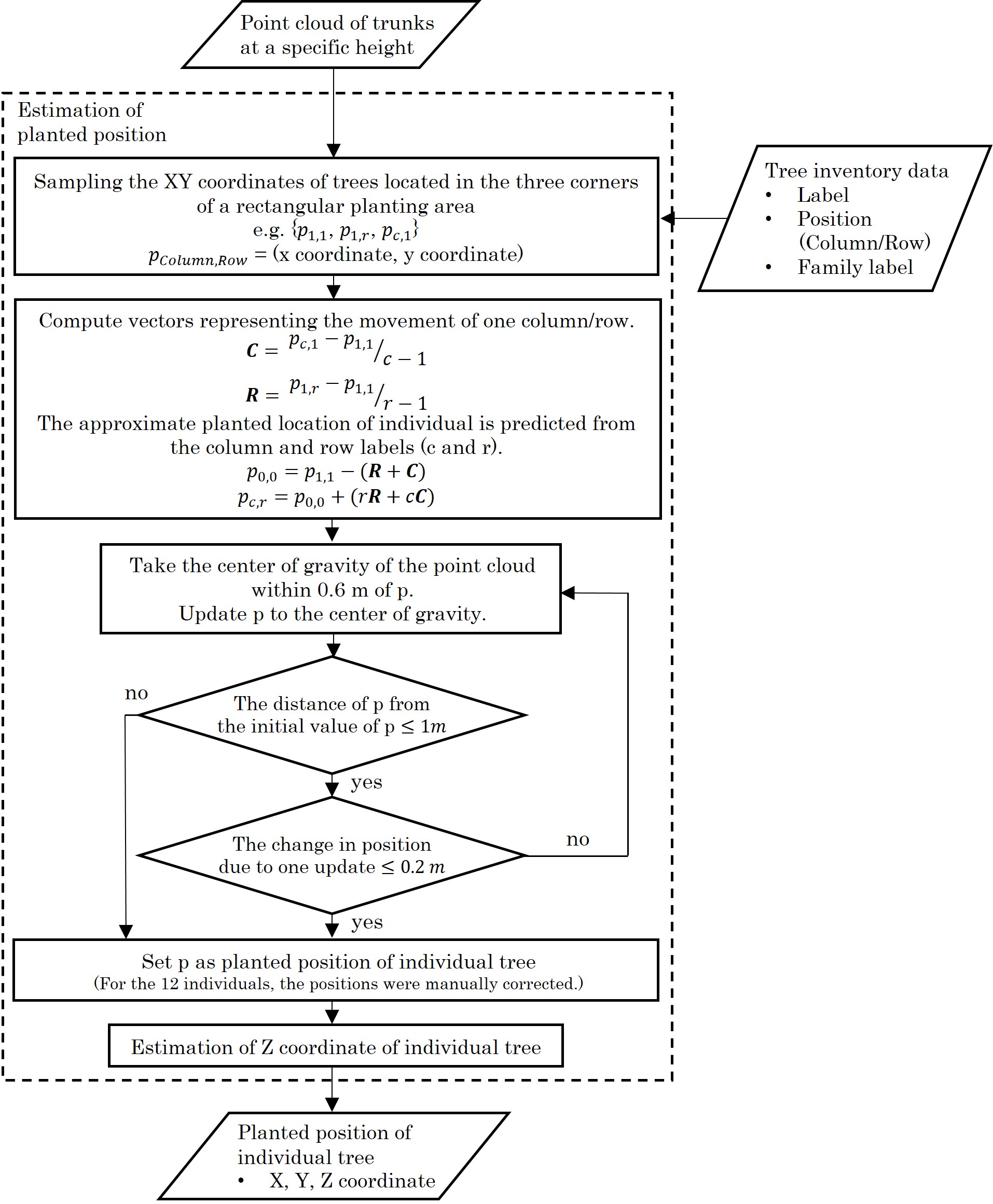

### Supplemental figure 3

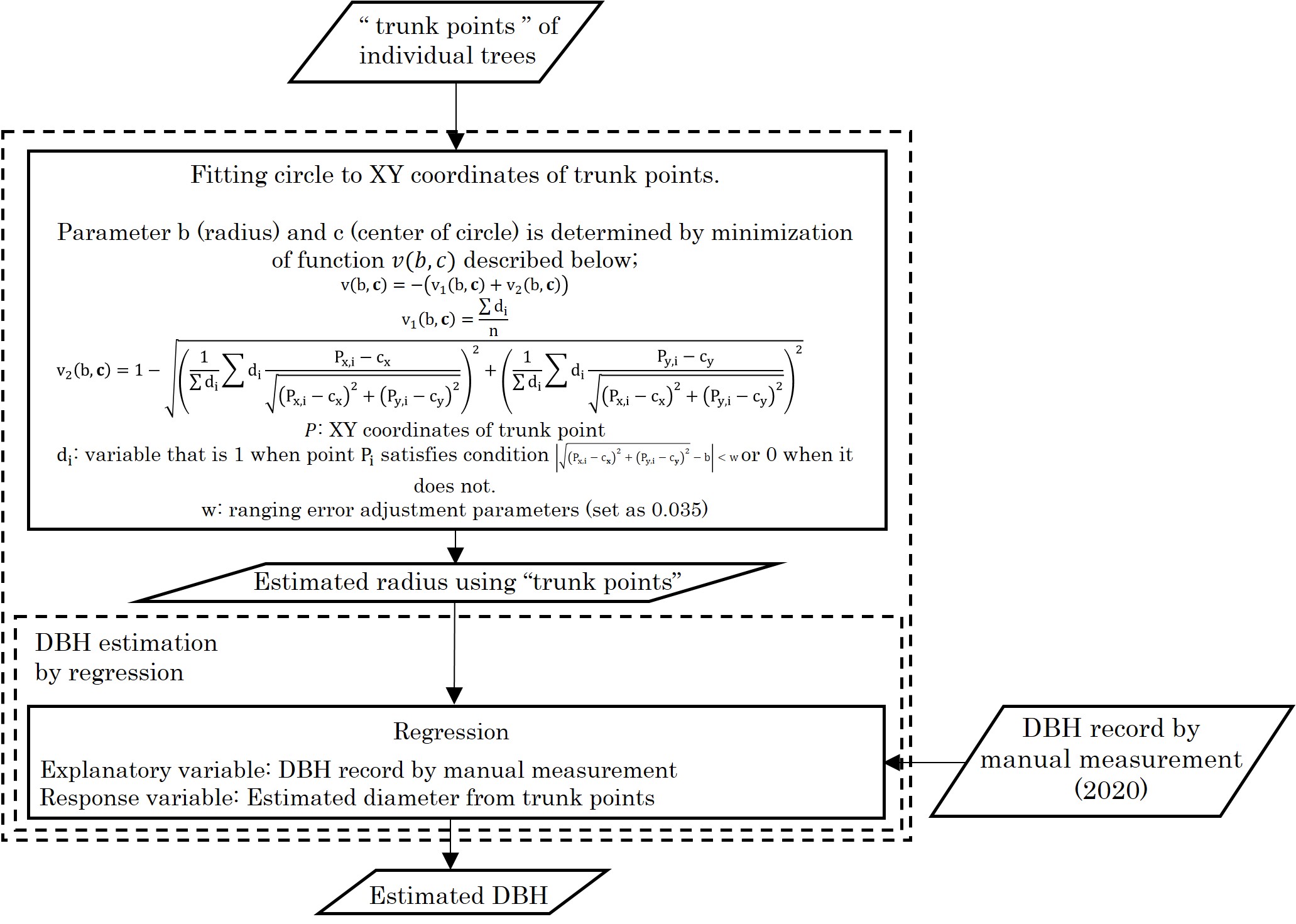

### Supplemental figure 4

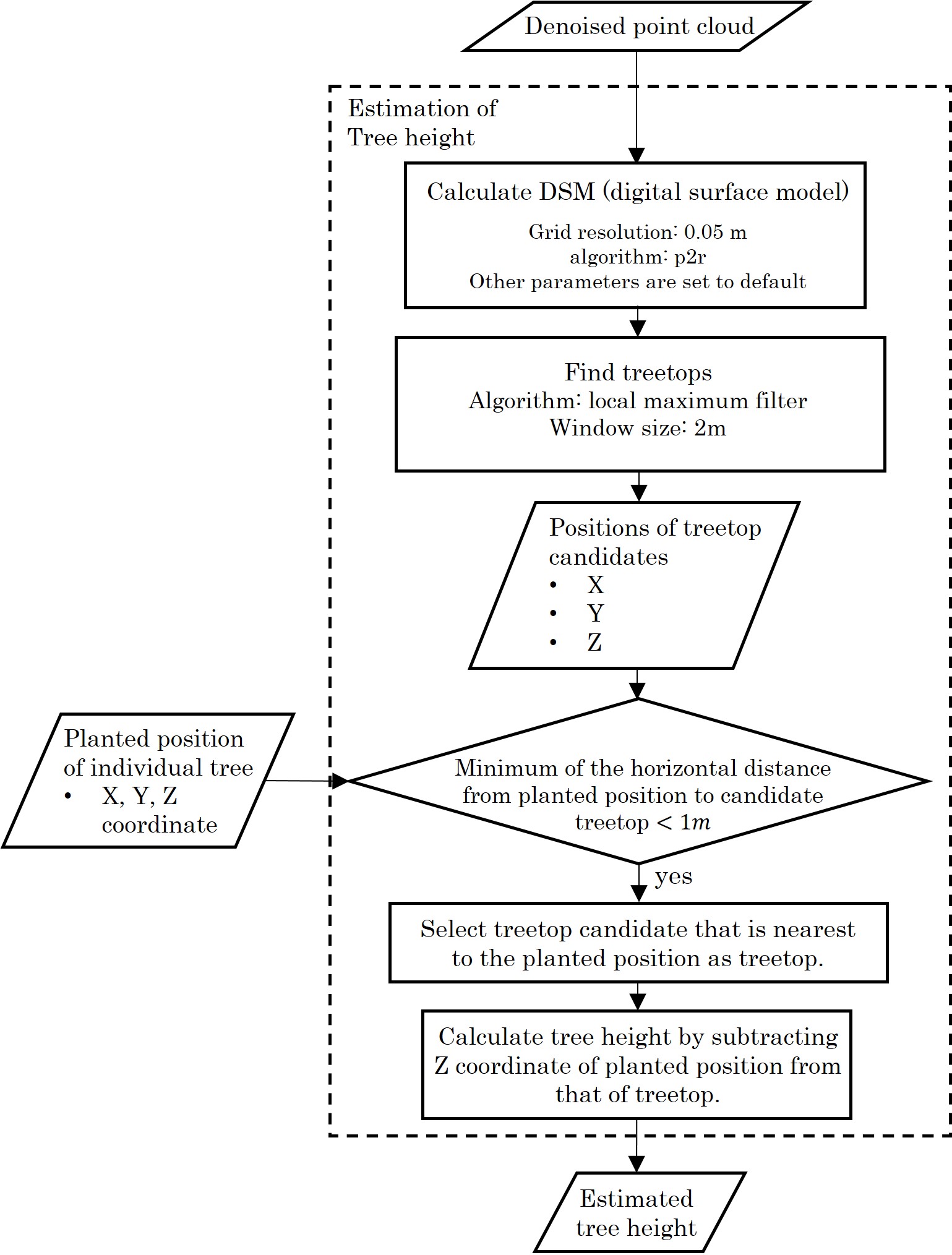
